# Optimized genome-wide CRISPR screening enables rapid engineering of growth-based phenotypes in *Yarrowia lipolytica*

**DOI:** 10.1101/2024.06.20.599746

**Authors:** Nicholas R. Robertson, Varun Trivedi, Brian Lupish, Adithya Ramesh, Yuna Aguilar, Anthony Arteaga, Alexander Nguyen, Sangcheon Lee, Chase Lenert-Mondou, Marcus Harland-Dunaway, Robert Jinkerson, Ian Wheeldon

## Abstract

CRISPR-Cas9 functional genomic screens uncover gene targets linked to various phenotypes for metabolic engineering with remarkable efficiency. However, these genome-wide screens face a number of design challenges, including variable guide RNA activity, ensuring sufficient genome coverage, and maintaining high transformation efficiencies to ensure full library representation. These challenges are prevalent in non-conventional yeast, many of which exhibit traits that are well suited to metabolic engineering and bioprocessing. To address these hurdles in the oleaginous yeast *Yarrowia lipolytica*, we designed a compact, high-activity genome-wide sgRNA library. The library was designed using DeepGuide, a sgRNA activity prediction algorithm, and a large dataset of ∼50,000 sgRNAs with known activity. Three guides per gene enables redundant targeting of 98.8% of genes in the genome in a library of 23,900 sgRNAs. We deployed the optimized library to uncover genes essential to the tolerance of acetate, a promising alternative carbon source, and various hydrocarbons present in many waste streams. Our screens yielded several gene knockouts that improve acetate tolerance on their own and as double knockouts in media containing acetate as the sole carbon source. Analysis of the hydrocarbon screens revealed genes related to fatty acid and alkane metabolism in *Y. lipolytica*. The optimized CRISPR gRNA library and its successful use in *Y. lipolytica* led to the discovery of alternative carbon source-related genes and provides a workflow for creating high-activity, compact genome-wide libraries for strain engineering.

**Highlights:**

1. Designed a compact, high activity CRISPR sgRNA knockout library for *Yarrowia lipolytica*.
2. Developed an efficient pipeline for discovering genes involved in alternative carbon-source utilization.
3. Identified single and double gene knockouts that improve growth on acetate.
4. Identified genes with improved fitness and essentiality for hydrocarbon growth.

## 1. Introduction

High throughput functional genomic screens are an essential tool for optimizing yeast production strains (Liu et al., 2015). By screening for the genotypes that underpin traits such as antimicrobial resistance (Kwak et al., 2011), ethanol resistance (Fujita et al., 2006), or morphological changes (Lupish et al., 2022), functional genomic screens can identify and deconvolute coding and regulatory sequences associated with specific cellular functions (Lian et al., 2019). Functional genome-wide screens have been especially effective tools for engineering industrially beneficial traits into yeast (Patterson et al., 2018). Yeasts such as *Saccharomyces cerevisiae* (Chen et al., 2016; Fujita et al., 2006), *Yarrowia lipolytica* (Jagtap et al., 2021; Lupish et al., 2022; Patterson et al., 2018; Ramesh et al., 2023), *Candida albicans* (Adames et al., 2019; Gervais et al., 2021; Wang et al., 2020), and *Komagataella phaffii* (Alva et al., 2021; Nishi et al., 2022; Tkachenko et al., 2023) have all undergone genome-wide screens to better understand industrially significant metabolic pathways. The outputs have led to promising improvements, such as increased stress tolerance (Crook et al., 2016; Fujita et al., 2006), improved protein production and secretion (Huang et al., 2015; Wang et al., 2019), and alternative nutrient utilization (Xu et al., 2019).

Many recent breakthroughs with functional genomic screens were enabled by CRISPR-editing methods, including CRISPR knockout (Horwitz et al., 2015), knock-in (Stovicek et al., 2015), CRISPRi (targeted gene inhibition (Momen-Roknabadi et al., 2020; Schwartz et al., 2017)), and CRISPRa (targeted gene activation (Schwartz et al., 2018) screens. These techniques provide broad genomic coverage and high specificity to identify gene targets in various host organism cellular functions. The key component of all CRISPR screens is the guide RNA (gRNA or sgRNA) library (Jakočiūnas et al., 2015), which may target either all of the genes in a host genome (Schwartz et al., 2019a), or a specific subset (Thompson et al., 2021). Once a library is assembled, CRISPR screens can be utilized to identify key genes in metabolic pathways or industrially relevant phenotypes (Jakočiūnas et al., 2015; Thorwall et al., 2020), or find gene expression changes that enhance pathway flux (Da Silva and Srikrishnan, 2012).

Functional CRISPR screens are categorized as either arrayed or pooled screens. Individually prepared mutants may be assembled into discrete or overlapping panels, and analyzed with an arrayed screen approach (Cachera et al., 2024). The outputs from arrayed screens are typically directly measurable phenotypes resulting from each mutant, such as production strain survival or accelerated growth rate (Garcia et al., 2021; Gutmann et al., 2021; Lupish et al., 2022). Arrayed screens are excellent for in-depth, quantitative analysis of specific genes predicted to affect phenotypes of interest (Liu et al., 2022). However, they are labor intensive to execute with larger libraries, as they require isolated strain preparations for each mutant to be tested (de Groot et al., 2018).

Alternatively, large, whole genome mutant libraries may be analyzed through a pooled screen (Bowman et al., 2020; Trivedi et al., 2023). Highly efficient transformation methods are used to generate a representative population of library cells, generating all possible single knockouts (or gene inhibitions/activations) of the given organism within a single culture volume (Schwartz et al., 2019). The pooled mutant library cells may then undergo phenotypic divergence when grown under a selection pressure (Smith et al., 2016). While individual genotypes cannot be directly observed or isolated in the pooled culture, deep sequencing can be used to find sgRNA prevalences and by extension, essential genes or genes involved in a phenotype of interest (Ramesh et al., 2023). Pooled genome-wide screens are an ideal high-throughput method for studying unknown relationships between genes, how those relationships affect desired phenotypes, and how those interactions affect metabolite production (Li et al., 2020). Still, challenges remain in the design and utilization of genome-wide sgRNA libraries. They must have maximized coverage of the host genome, ideally with highly efficient guide activities. Guide RNA expression characteristics must also be optimized to ensure sufficient genome coverage (Dalvie et al., 2020). To overcome these challenges, techniques are required to better optimize CRISPR gRNA libraries.

In this work, we design an optimized sgRNA library for CRISPR genome-wide pooled screens in *Y. lipolytica*. We then validated our library for use in growth-based functional genomic screens to improve growth on non-glucose carbon sources, including acetate, fatty acids, and hydrocarbons. To optimize our library, we strategically reduced the size of a previously established *Y. lipolytica* sgRNA knockout library by half, while maintaining coverage of the host genome. In addition to existing experimentally validated sgRNAs, we also incorporated new sgRNAs predicted by DeepGuide (Baisya et al., 2022). To identify phenotypes of interest, we screened *Y. lipolytica* harboring both the sgRNA library and integrated Cas9 on a variety of carbon sources. From our hydrocarbon screens, we found significant overlap in genes essential for growth between similar hydrocarbons. We experimentally validated our acetate hits and found that knockouts of our top hits shortened lag phase and increased growth with acetate as the sole carbon source. This *Y. lipolytica* optimized CRISPR gRNA library provides a workflow for designing efficient libraries for functional genomic screening of metabolic engineering traits.

## 2. Results and Discussion

### 2.1. Compact CRISPR-Cas9 library contains high-activity sgRNAs

Iterating on our previously created 6-fold coverage library, we sought to create a minimally sized, high activity “v2” library. The high activity gives more confidence in the data and the smaller size reduces the transformation burden to achieve high coverage libraries. We designed a 3-fold coverage CRISPR-Cas9 library to target all protein-coding sequences in the PO1f strain of *Y. lipolytica*. The library consists of a combination of high-activity sgRNAs from an existing *Y. lipolytica* CRISPR-Cas9 library designed by (Schwartz et al., 2019b) (denoted as “v1”) and sgRNAs predicted to be of high activity by an activity prediction tool, DeepGuide (Baisya et al., 2022) (**Fig. 1a,b**).

**Figure 1.**
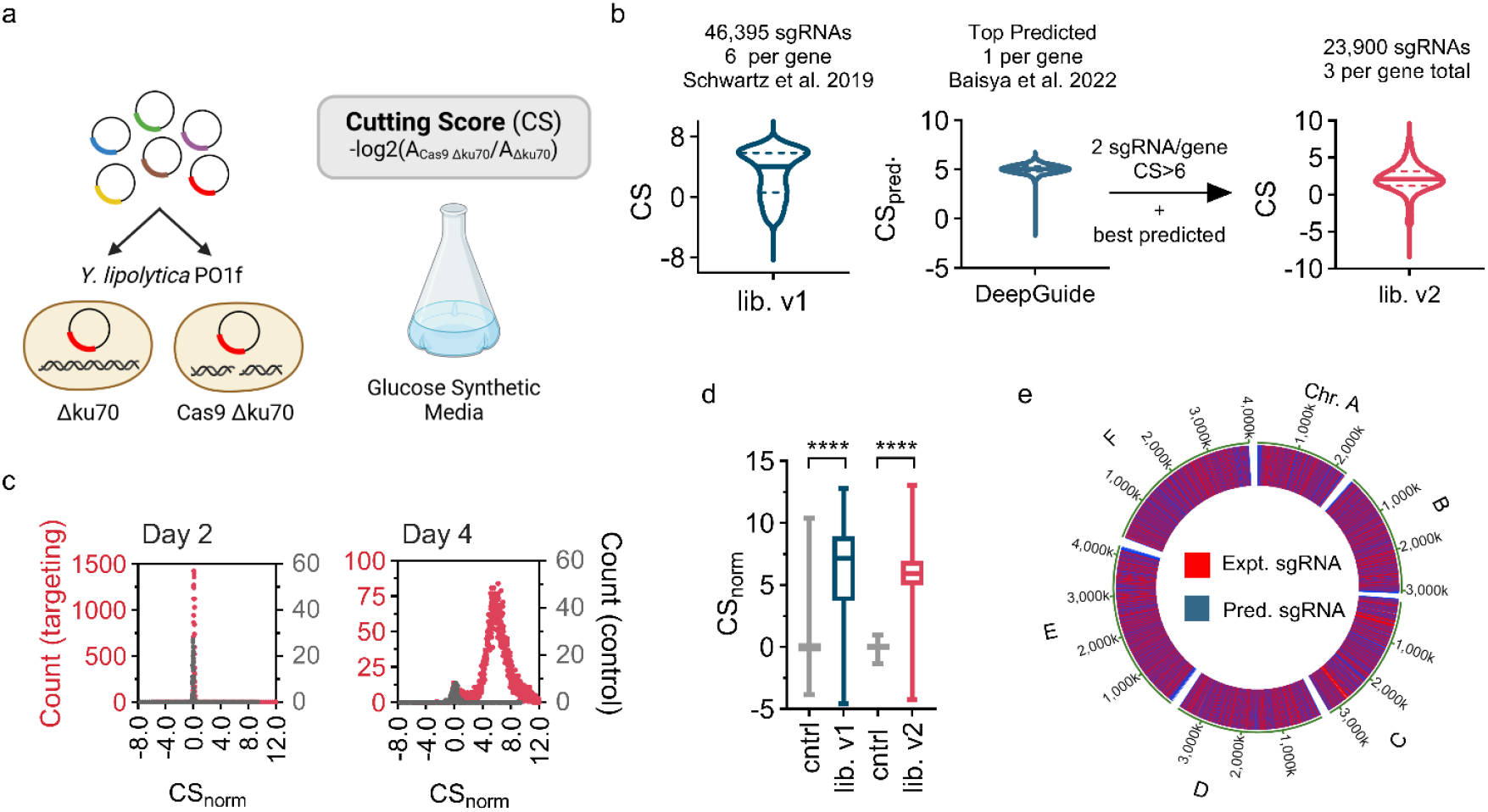
Optimized CRISPR library generation pipeline and characterization. (a) Cutting score (CS) is calculated as the depletion of a guide in the Cas9 *ku70* knockout strain compared to the *ku70* knockout control. A guide which cuts well in the Cas9 *ku70* knockout strain will become depleted as cells die or have a serious growth penalty. (b) The two best cutting guides from lib. v1 were used in addition with the top predicted guide from DeepGuide to generate lib. v2. (c) Distributions of targeting (red) and non-targeting (gray) sgRNAs after two and four days of growth post transformation. After four days of growth with one subculture on day 2, the high activity sgRNAs diverge from the non-targeting population. (d) Both lib. v1 and v2 contain active guides, however, v2 contains more guides with high activity. (e) DeepGuide predicted and lib. v1 guides are distributed randomly throughout the genome.

Growth screens previously conducted in the *Y. lipolytica* PO1f strain using library v1 defined a metric called cutting score (CS) for each sgRNA — a measure of its cutting efficiency — by comparing sgRNA abundance in the PO1f Cas9 Δ*KU70* strain to that in the control strain devoid of Cas9 (Schwartz et al., 2019b). Guides that make a cut in the experimental strain become depleted as cells die since disruption of *KU70* suppresses non-homologous end joining, the dominant DNA repair mechanism in *Y. lipolytica* (Schwartz et al., 2017), whereas guides that do not make a cut become enriched. By comparing the abundance of each guide in the experimental strain to the abundance of the same guide in the control strain, CS can be calculated. Activity predictions from DeepGuide were generated by training the tool using sgRNA sequences from library v1 and MNase-seq-derived nucleosome occupancy data for *Y. lipolytica* PO1f as input data and the CS of every sgRNA as output. The result is a convolutional neural network model that accurately predicts high activity CRISPR-Cas9 sgRNAs (Baisya et al., 2022). In order to pick sgRNAs for library v2, library v1 was first filtered to only retain sgRNAs having a CS greater than 4.0 (a value close to the optimum CS threshold for high-activity sgRNAs previously determined by (Ramesh et al., 2023). Library v2 was thus designed to contain the top two guides with CS > 4.0 for each gene from the filtered library v1, and a guide with the highest DeepGuide-predicted CS for that gene. If, for a gene, less than 2 sgRNAs from library v1 had CS > 4.0, appropriate numbers of sgRNAs with high DeepGuide-predicted CS were included so as to maintain a library coverage of 3 (see Materials and Methods). The final library consists of three sgRNAs per gene targeting ∼99.8% of all genes (**Fig. S1a**). Moreover, next generation sequencing of the cloned, untransformed library revealed a tight normal distribution of sgRNA abundances, indicating a near-uniform and symmetrical (median ≈ mean) distribution of guides in the library (**Fig. S1b**). sgRNA cutting scores for library v2 were generated using the same screening framework as that used for obtaining CS of library v1 guides. The CS for all guides in the v2 library are provided in **Supplementary File 1**.

In addition to gene-targeting guides, library v2 also consists of 360 non-targeting sgRNAs (∼1.5% of the total library) that serve as negative controls. The normalized CS (*i.e.*, the difference between raw CS of a guide and the average CS of non-targeting guides) distributions of targeting and non-targeting sgRNAs nearly overlap on day 2 (avg. CS_norm, targeting_ = 0.12±0.09 and avg. CS_norm, non-targeting_ = 0±0.06) due to no observable effect of CRISPR-induced double stranded breaks on cell growth. On day 4, however, the two populations are well separated (avg. CS_norm, targeting_ = 5.96±2.04 and avg. CS_norm, non-targeting_ = 0±0.34), indicating different activity profiles of targeting and non-targeting guides (**Fig. 1c**). The difference between targeting and non-targeting populations is statistically higher for library v2 (one-tailed unpaired t-test p < 0.0001, t-score = 58.81) compared to v1 (one-tailed unpaired t-test p < 0.0001, t-score = 43.49), providing evidence that library v2 is a higher-activity library compared to library v1 (**Fig. 1d**). Also of note is the genomic loci of the experimental and DeepGuide-predicted sgRNAs used to construct library v2, which are evenly distributed across the genome (**Fig. 1e**). These data show that we built a compact, high activity library that can be used in functional genomic screens in *Y. lipolytica*. The small size enables high library coverage due to lower transformation burden and the high activity allows near complete assurance of genomic coverage.

### 2.2. Acetate tolerance screens identify gene targets for improved growth

With the optimized genome-wide CRISPR library in hand, we next set out to identify genes essential for growth on acetate. Functional genomic screens were conducted using minimal media supplemented with three different carbon sources — glucose (2% (w/v)) as a control condition, and low and high acetate (250 and 500 mM, respectively) as experimental conditions (**Fig. 2a**). These screens allowed us to calculate a fitness score (FS) for every sgRNA in each media condition. A high FS value indicates that knockout of that gene gives a fitness advantage, provided a cut happens. Conversely, low FS values indicate a loss of cell fitness due to disruption of the target gene. FS is computed as the log_2_-ratio of sgRNA abundance in the PO1f Cas9 strain to that in the PO1f control strain. We used acCRISPR, an analysis pipeline for CRISPR screens that correct screening outcomes based on the activity of each guide used in the screen (Ramesh et al., 2023) to identify essential genes in glucose, and high and low FS value genes in the two acetate conditions. The analysis pipeline identified 1580 essential genes for growth with glucose as the sole carbon source, as well as 868 and 901 genes that reduce or improve growth on acetate, respectively (**Fig. 2b**).

**Figure 2.**
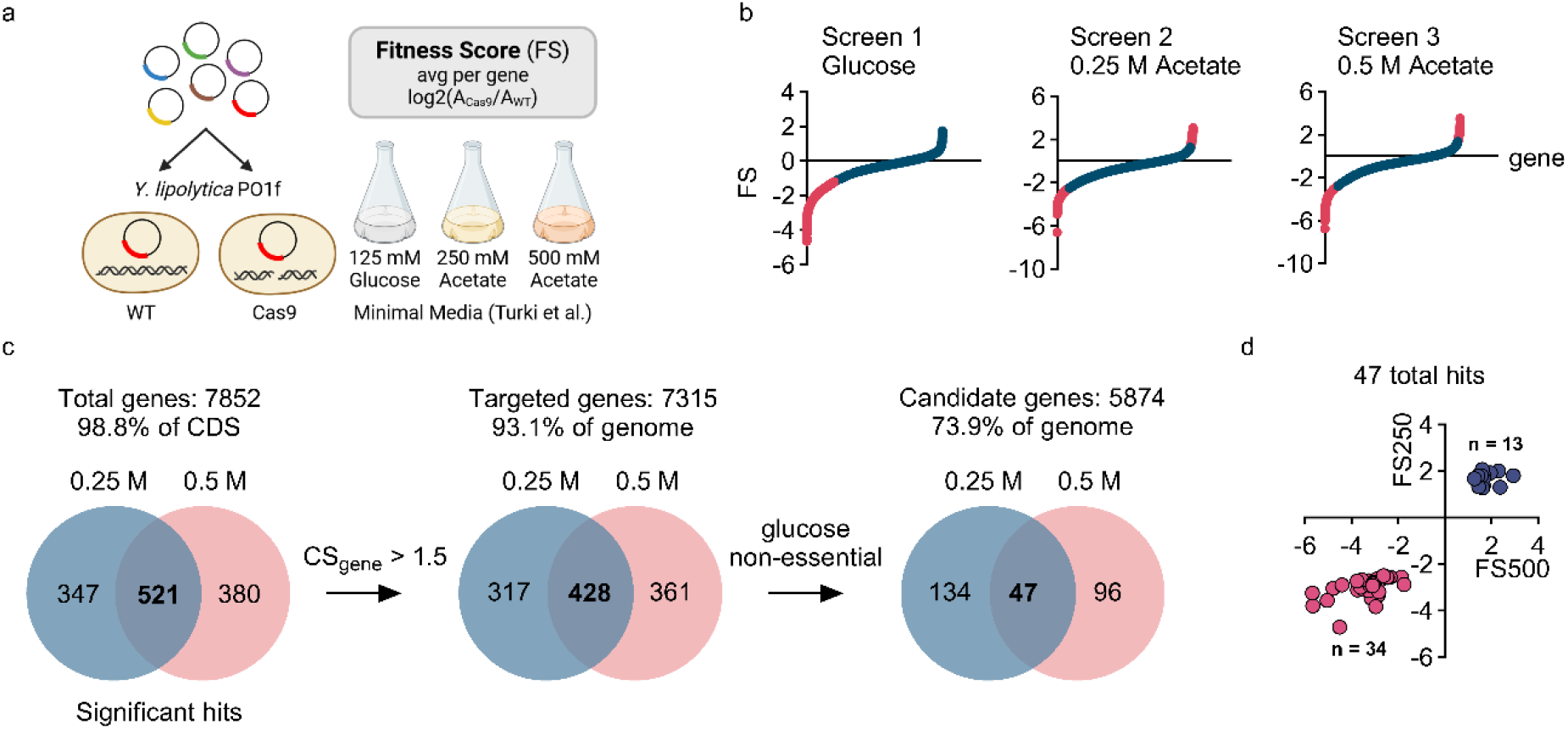
Growth-based functional genomic CRISPR screening with acetate as the sole carbon source. (a) Growth screens were conducted using *Y. lipolytica* PO1f and a strain containing Cas9 as control and treatment strains respectively in three media conditions - 125 mM glucose, 250 mM and 500 mM acetate. Fitness score (FS) is calculated as the average enrichment of guides targeting a gene in the Cas9 strain compared to PO1f. Genes with a higher FS indicate that the knockout gave a fitness advantage. (b) Corrected FS values for all genes for the glucose and acetate screens, as determined from the acCRISPR pipeline. Pink portion of the curve represents essential genes for glucose, and high and low FS genes for the two acetate conditions. (c) Venn diagrams representing gene hits in low and high acetate conditions after each filtering step. Low-activity guides and glucose essential genes were removed to condense the pool of hits. (d) Top low and high FS hits in both acetate conditions.

Despite lib. v2 containing mostly high activity cutters, we wanted to be sure that very few false positive hits would arise at the end of our screening pipeline. To do this, we used guides with high activity (CS > 1.5) for calling hits (**Fig. 2c**). This reduced the size of the common significant gene set between the two acetate conditions to 428 (13 with high FS values and 415 with low FS values). Any double stranded break that occurs in a cell causes at least a moderate fitness effect (Schwartz et al., 2019). Additionally, there are likely very few gene knockouts that improve tolerance to acetate in *Y. lipolytica*. These reasons may explain why we demonstrated an 86% loss of positive FS hits when we removed poor targeting guides. These false positives represent control strain cells that did not have a fitness disadvantage of a double stranded break. Next, we wanted to exclude any significant genes that reduce fitness in glucose media. We reasoned that the most relevant hits should be solely due to acetate fitness effects and not to overall fitness. Glucose viability is also important in upstream applications for making further strain modifications and for downstream applications if glucose supplementation is needed for an increase in intracellular reducing power. As a result, we excluded genes that were found to be essential in glucose from the significant gene lists for acetate, further condensing the common significant gene set to 47 genes. Thirteen of the 47 genes had positive FS values while the remaining 34 genes had FS values lower than 0 (**Fig. 2d**).

### 2.3. Acetate screen hits shorten lag phase growth

To validate the screening hits, we focused on the top 13 positive FS hits identified in the low and high acetate screening conditions, as knocking out these genes should improve growth on acetate (**Fig. 3a**). To do this, we transformed PO1f with a plasmid containing Cas9 and the relevant guide. We obtained 12 of the 13 top positive FS knockouts; functional gene disruptions were confirmed with colony PCR and sanger sequencing. Our top hit, D05956g, encodes SFL1, a flocculation suppressant gene. When knocked out, this gene causes *Y. lipolytica* to flocculate (**Fig. S2**). This mutation could provide a fitness advantage in the context of our screen as it allows cells to clump together at the bottom of the flask and avoid the high osmotic stress of being a suspended single cell. The flocculation behavior makes growth measurements challenging and thus the D05956g knockout was excluded from further analysis.

**Figure 3.**
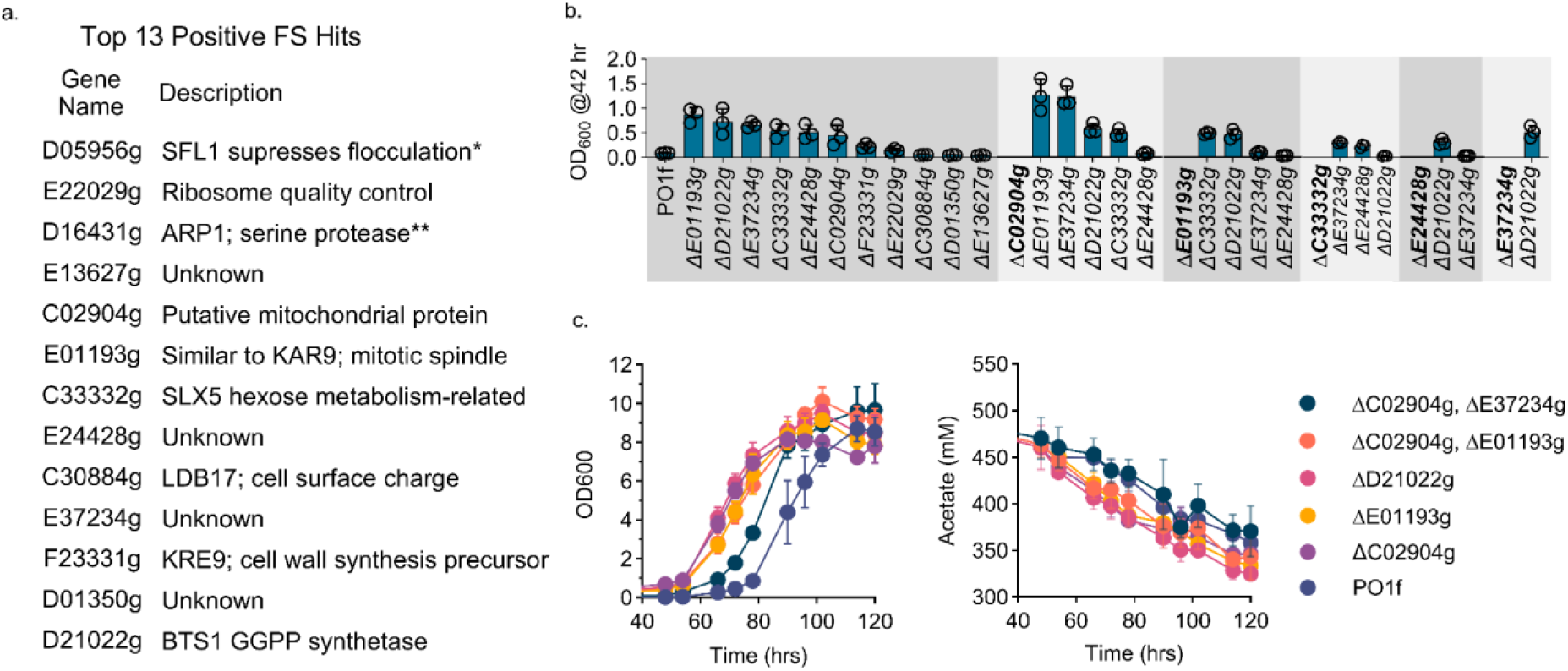
Validating acetate metabolism hits. (a) Thirteen positive FS value hits in descending order of high acetate FS. (b) Single (left gray box) and double knockout (5 gray boxes on the right) mutants tested in 96-well 1 mL wells containing 500 mM acetate in minimal media (30 °C, 42 hrs, 1000 RPM). Bars represent the mean of three biological replicates, error bars represent the standard deviation. Data points are shown for each replicate. (c) Growth curve and acetate depletion for top three single knockouts and top two double knockouts grown in 250 mL shake flasks with 500 mM acetate in minimal media (30 °C, 220 RPM). Acetate depletion measured with HPLC. Data points represent the mean of three biological replicates, error bars represent the standard deviation.

For the remaining 11 knockouts, we validated their phenotype in growth assays that mimic the screening conditions, growth in minimal media with 500 mM acetate as the sole carbon source. Under these growth conditions we observed that at 42 hours, *Y. lipolytica* PO1f begins to break out of the lag phase and enter the exponential growth phase. This time point gave us the most resolution and repeatability to confirm the hits. Of the 11 characterized knockouts, 8 grew significantly (p<0.05) better than the control after correcting for multiple comparisons, including E01193g, D21022g, E37234g, C33332g, E24428g, C02904g, F23331g, and E22029g (**Fig. 3b**). We next sought to explore if higher order mutations would further improve *Y. lipolytica* growth in acetate media. To test the effect of stacked mutations, we chose the top 6 knockouts (E01193g, D21022g, E37234g, C33332g, E24428g, and C02904g) and tested all 15 double knockout permutations. Only two double knockouts grew better than our top single knockout, possibly due to a combined reduction in general cell fitness in the higher order mutants.

We followed up the preliminary hit validation with more comprehensive characterization of cell growth and quantification of acetate depletion. The top two single and double knockouts were selected as well as C02904g due to the commonality between the top two double knockout hits (**Fig. 3c**). Limited overall growth and incomplete acetate consumption may be explained by micronutrient or amino acid depletion or pH change caused by consumption of acetate while leaving behind sodium ions (**Fig. S3a**). Roughly 25% of acetate was converted to dry cell weight (DCW) (**Fig. S3b**). Despite this, all hits grow better and consume more acetate than the PO1f control strain. In addition, lag phase was reduced by 24 hours in our most potent knockout strains. These data provide strong validation of our top positive FS hits as mutations that lead to the improvement of acetate metabolism in *Y. lipolytica*.

Our top four hits to improve growth were E37234g (unknown function), E01193g (KAR9, which plays a role in mitotic spindle positioning in *S. cerevisiae* (Tirnauer et al., 2000)), C02904g (FMP42, putative mitochondrial protein in *S. cerevisiae* (Imamura et al., 2015)), and D21022g (GGPP synthetase homolog (Saikia et al., 2009, Morton et al., 2011)). GGPP synthetase converts farnesyl pyrophosphate (FPP), the precursor to steroids and N-glycans, to GGPP, a precursor to terpenoids and other metabolites (Chen et al., 2024, Liu et al., 2014). Previous work has shown that deletion of this gene improves oxidative stress response in *N. crassa* (Sun et al., 2019). One possibility is that knockout of GGPP synthetase increases intracellular FPP and decreases GGPP, which may reduce oxidative stress from endogenous terpenoid production and increase cell wall integrity by improved flux to membrane essential compounds through FPP.

### 2.4. Hydrocarbon screening reveals genes necessary for fatty acid metabolism and transport

Given the success of the genome-wide CRISPR approach in identifying genes that improve growth on acetate, we next sought to uncover genes related to hydrocarbon and fatty acid metabolism, substrates on which *Y. lipolytica* has a demonstrated ability to grow (Fickers et al., 2005; Try et al., 2018). Dodecane, 1-dodecene, oleic acid, and margaric acid were chosen as substrates for their diversity in polarity and hydrogenation. Preliminary experiments showed that solid media produced more consistent cell growth than liquid cultures, as such this format was used for functional genomic screening. Glucose as a sole carbon source was also screened to control for any putative hits that were solely due to the transition to solid media (**Fig. 4a**).

**Figure 4.**
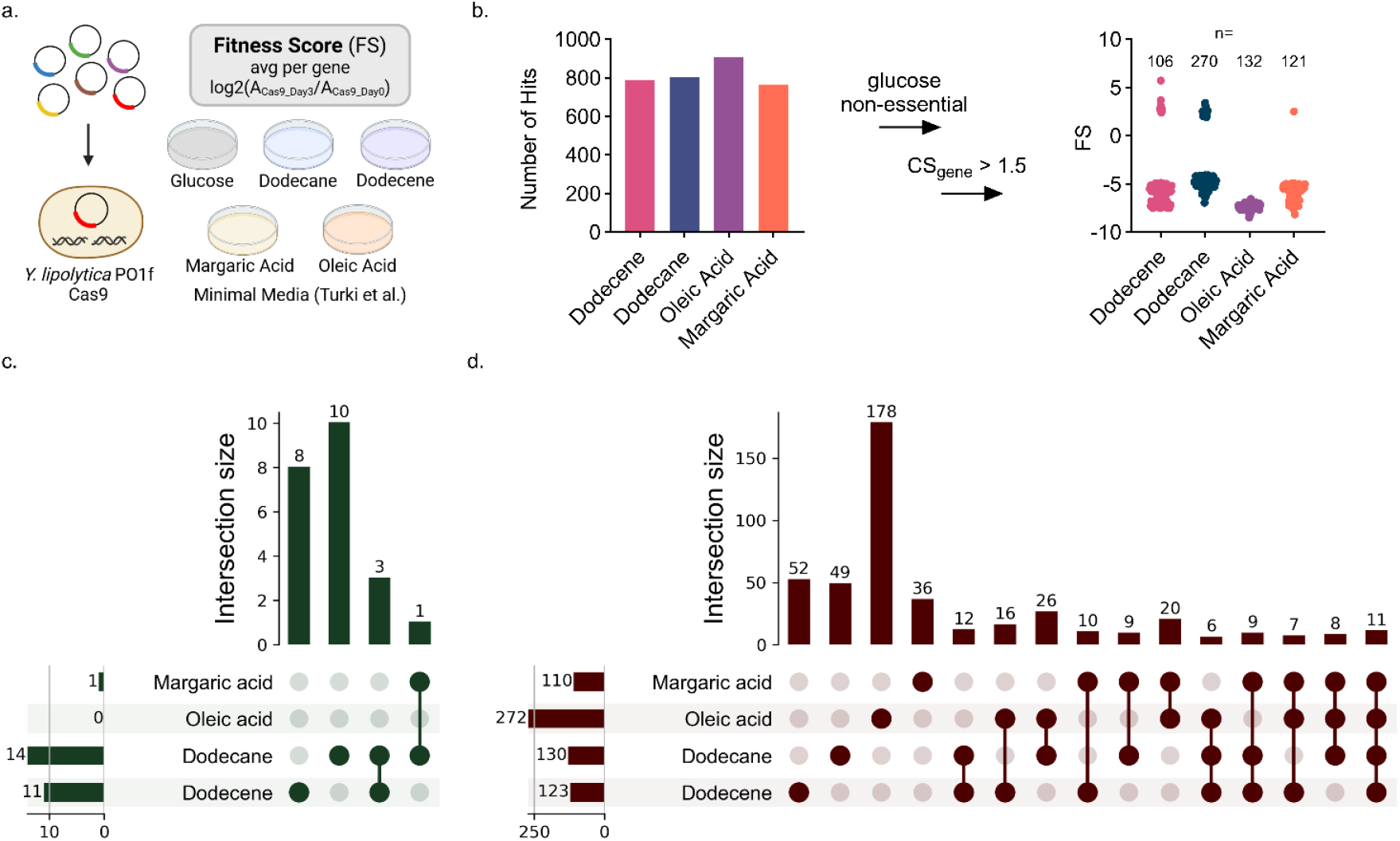
Growth-based functional genomic CRISPR screening with hydrocarbon substrates. (a) Growth screen, carbon sources screened and fitness score (FS). Genes with a higher FS indicate that the knockout gave a fitness advantage. (b) Filtering of gene hits obtained for each hydrocarbon condition; gene hits with poor cutting guides and genes essential for growth on glucose were removed to obtain high confidence hits. (c,d) UpSet plot analysis for hits in all hydrocarbon conditions. Hits with a fitness advantage are shown in (c), while essential hits are shown in (d). The values associated with each substrate indicate the number of gene hits that were identified for each substrate. Screen growth conditions were 55.5 mM dodecane solid media grown for 48 h, 55.5 mM 1-dodecene solid media grown for 48 h, 37 mM oleic acid solid media grown for 48 h, and 39.2 mM margaric acid solid media grown for 72 h, all at 30 C in a static incubator, with two screen replicates of each condition, with each replicate consisting of seven 100 mm plates.

Hits were identified in a similar manner to the acetate screen: the acCRISPR analysis pipeline was used to determine FS values from next generation sequencing read counts of the sgRNAs remaining at the end of the screen; genes with fitness advantage or essentiality were called as hits; hits from genes with low CS guides were removed from the hit pool to avoid false positives; and, genes found to be essential in the glucose liquid culture condition were removed from the hit list. In total 478 high confidence hits were called across the four substrates tested, with 28 gene knockouts with a fitness advantage, and 449 genes essential for growth on the substrates (**Fig. 4b**).

Upset plot analysis is a visual means of identifying hits that are common across the various conditions (**Fig. 4c,d**). When applied to our screens, this analysis revealed that three hits with positive FS values were called in both the dodecane and dodecene screens, each of which is involved with membrane transport of lipids or carbohydrates (**Fig. 4c**). The genes common to the dodecane and dodecene screens were a mixture of stress response, endomembrane trafficking, and pro-hyphal membrane and cell wall maintenance genes. There were no positive FS hits for oleic acid and only one for margaric acid (C06486g, a knockout of a transcription factor with homology to *S.pombe ADN1*, believed to promote invasive growth and cell adhesion), which was also found in the dodecane screen. In addition to these hits, ten positive FS hits were identified for dodecane and eight for dodecene (see **Supp. File 2** for a list of all genes in the upset plots). Additional analysis of screen hits with positive FS is shown in **Fig. 5**. With respect to the essential genes (*i.e.*, those with low FS values), the overlap between carbon sources was more comprehensive (**Fig. 4d**). Eleven genes were found to be common across all screens, another 30 were common to three of four screens and 93 were common to two of four screens. Functional analysis of these genes, as well as hits with a fitness advantage, are presented in **Fig. 5**.

**Figure 5.**
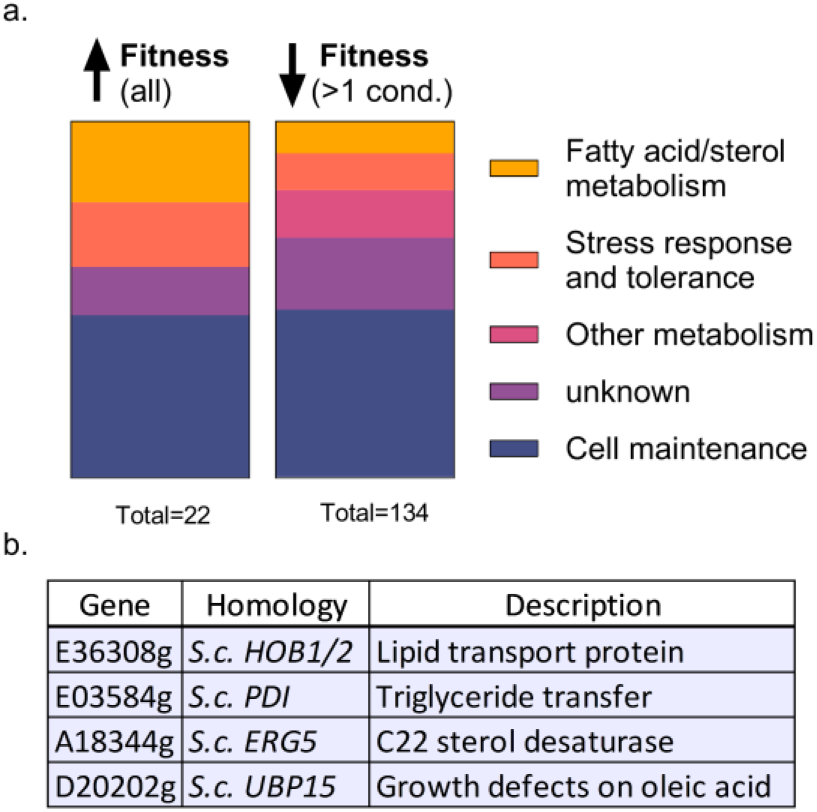
Biological function of top hydrocarbon screening hits. (a) Hit distribution by biological function: Fatty acid/sterol metabolism, stress response and tolerance, cell maintenance, and unknown. The left bar includes all positive FS gene hits in all hydrocarbon screens (n=22). The right bar includes all essential gene hits called in two or more screens (n=134). (b) Fatty acid metabolism-related gene hits that improve fitness in at least one hydrocarbon condition.

Our screen revealed 134 total genes that reduce fitness in at least two hydrocarbon/fatty acid conditions (**Fig. 5a**). Twelve hits that reduced fitness were directly related to hydrocarbon metabolism. One of these twelve was C03415, a hit identified in all four hydrocarbon screens and known to be essential for fatty acid metabolism in *Aspergillus nidulans* (*acuH* homolog) (De Lucas et al., 1999). Another 18 hits were related to metabolic pathways, including endomembrane trafficking genes. These include E23880g, an essential gene for all four conditions which is homologous to *S.cerevisiae VPS29*, a membrane-bound vesicle reverse-transport protein (Seaman et al., 1998)), and D14097g that was found to be essential in three of the four screens. D14097g is homologous to *S.cerevisiae VPS53*, which is responsible for retrograde sorting in the golgi apparatus (Conibear and Stevens, 2000). We expected hits such as these two, as there is experimental evidence that these genes are involved with endomembrane trafficking that affect alkane metabolism (Fukuda, 2013). Other hits include the previously mentioned *ADN1* homologue, and E07121g found in our dodecane hits (homologous to *PMT4* in *C. albicans* which plays a role in invasive hyphal growth (Lengeler et al., 2008)). We expected a fitness advantage from deletions of pro-hyphal genes like these since there is a known inverse relationship between hyphal morphology and efficient hydrocarbon metabolism (Palande et al., 2014). In addition to the hydrocarbon and metabolic pathway hits, there were 14 hits related to stress response/tolerance genes, a class of genes that are expected to be essential for growth under these conditions. Last, there were 63 hits related to cell maintenance functions, which we anticipated as they are essential for maintaining cell viability when hydrocarbons are the only available carbon source. Taken together, these hits reveal metabolic pathways and cellular functions that are essential for survival in hydrocarbons and fatty acids.

In addition to the hydrocarbon essential genes, we identified 22 genes that produced positive FS values in the hydrocarbon and fatty acid screens and are beneficial to growth on one or more of the hydrocarbon substrates. Similar to the essential gene hits, roughly half of these hits were related to cell maintenance. This group includes E30725g, identified in the 1-dodecane screen and a homolog of *MNN10* in *S. cerevisiae,* and is related to cell membrane/wall maintenance ((Jungmann et al., 1999; Klein et al., 2002; Sacristán-Reviriego et al., 2014)). C06486g, an *ADN1* homolog in *S. pombe*, was found in both margaric acid and dodecane (Dodgson et al., 2009). This gene is involved in cell adhesion and hyphal regulation, a trait known to affect hydrocarbon metabolism ((Palande et al., 2014). Five genes related to stress response were also found to improve growth on hydrocarbons. Specifically, C14369g (homologous to various *Candida* species, related to azole drug resistance ((Costa et al., 2013; Dong et al., 2021))) and F01386g (homologous to *MSG5* in *S. cerevisiae,* involved in cell signaling (Jungmann et al., 1999; Klein et al., 2002; Sacristán-Reviriego et al., 2014))) were found in the dodecane and 1-dodecene screens, respectively. We uncovered four hits directly related to improved tolerance to hydrocarbons (**Fig. 5b**). The *HOB1/2* (E36308g) homolog in *S. cerevisiae* was present in both dodecane and 1-dodecene and is a lipid transport protein (Castro et al., 2022). The remaining three genes were only found in 1-dodecene and include a triglyceride transfer protein (E03584g, PDI homolog in *S. cerevisiae* (Gauss et al., 2011), a C22 hydrocarbon sterol desaturase (A18344g, *ERG5* homolog in *S. cerevisiae* (Skaggs et al., 1996), and an *S. cerevisiae* UBP15 homolog found to cause growth defects on oleic acid (Debelyy et al., 2011) (**Fig. 5b**). In both fitness improvement and essential gene hits, our screen outputs demonstrate how genome-wide CRISPR knockout screens are a vital component of efficient strain engineering.

## 3. Conclusions

Genome-wide CRISPR screening in industrial hosts like *Y. lipolytica* enables rapid engineering of growth-based phenotypes. We previously developed a genome-wide CRISPR-Cas9 knockout library for use in Yarrowia that can be used in functional genetic screens and in rapid design-build-test-learn cycles for strain engineering. The first version of this screening tool contained sgRNAs with a wide range of activity and needed 6-fold coverage to cover nearly all the genes in the genome. Here, we create a new optimized, high activity sgRNA library that is compact in size. This library required only 3-fold genome coverage to target 98.8% of genes in the genome. By creating a library one-half the size, we reduce the burden of generating large libraries that require high transformation efficiency to obtain. Quantifying the activity of each guide in the library revealed that all but a handful of guides are highly active. A tight, normal distribution of guide activity enables high accuracy screening as all (or nearly all) guides can be used to identify gene hits. This optimized CRISPR-Cas9 screening tool enabled us to conduct high throughput strain engineering experiments for growth on various alternative carbon sources, including acetate, dodecane, 1-dodecene, oleic acid, and margaric acid. The acetate screens identified several single and double mutant strains with shortened culture times when consuming acetate as the sole carbon source. The hydrocarbon screens revealed 22 gene targets for improving growth and a set of 134 genes essential for fatty acids and long chain hydrocarbon metabolism. Many of these hits have unknown function or are gene targets for metabolic engineering. Our compact, high activity CRISPR-Cas9 library enables growth-based screens for enhancing carbon source utilization and promises to enable a wide range of screens for improving other growth-based phenotypes.

## 4. Materials and Methods

### 4.1. Microbial strains and culturing

All strains used in this work are presented in **Table S1**. *Yarrowia lipolytica* PO1f (MatA, leu2-270, ura3-302, xpr2-322, axp-2) is the parent for all mutants used in this work. Unless otherwise noted, all yeast culture growth was carried out in 14 mL polypropylene tubes or 250 mL baffled flasks, with incubator conditions of 30 °C and 220 RPM. Under non-selective conditions, *Y. lipolytica* was grown in YPD (1% Bacto yeast extract, 2% Bacto peptone, 2% glucose). Cells transformed with sgRNA-expressing plasmids were initially propagated in synthetic defined media deficient in leucine (SD-leu; 0.67% Difco yeast nitrogen base without amino acids, 0.069% CSM-leu (Sunrise Science, San Diego, CA), and 2% glucose) for two days to allow for genome edits to occur. All plasmid constructions and propagations were conducted in *Escherichia coli* TOP10. *E. coli* cultures grown in Luria-Bertani (LB) broth with 100 mg/L ampicillin at 37 °C in 14 mL polypropylene tubes, at 220 RPM. Plasmids were isolated from *E. coli* cultures using the Zymo Research Plasmid Miniprep Kit II.

### 4.2. Plasmid construction

All plasmids and primers used in this work are listed in **Table S2,3**. The plasmids used to knock out genes in *Y. lipolytica* PO1f were constructed by ordering the corresponding sgRNA as a primer (**Table S4**) with 20 bp homology up- and downstream of the AvrII cutsite in the pSC012 plasmid. 60 bp top and bottom strands were ordered and annealed together. The annealed strand and digested plasmid were assembled using Gibson Assembly in a 10:1 molar ratio (insert:vector). The assembly product was then transformed directly into electrocompetent *E. coli* TOP10 cells to eventually be propagated and harvested by Miniprep.

### 4.3. *Y. lipolytica* CRISPR knockouts

A plasmid containing Cas9 and the appropriate sgRNA (pSC012) for the desired gene knockout was transformed into *Y. lipolytica* using a protocol described by Chen et al. (Chen et al., 1997). In short, a single colony of the background strain of interest was grown in 2 mL of YPD liquid culture in a 14 mL culture tube at 30 °C with shaking at 220 RPM for 22-24 hours (final OD ∼30). 300 µL of culture (∼10^8^ cells) were pelleted by centrifugation at 4,000g for 2 minutes and then resuspended in 300 µL of transformation buffer. The transformation buffer contains a final concentration of 45% PEG 4000, 0.1M Lithium Acetate and 100 mM Dithiothreitol. Then, 500 ng of plasmid DNA was added followed by 8 µL 10 mg/L ssDNA (Agilent). The reaction mix was vortexed thoroughly and then incubated for 1 hour at 39 °C. 1 mL of water was added and then the cells were pelleted and inoculated into 2 mL SD-ura selective liquid media. After 3 days, the cells were plated at a 10^−6^ dilution on YPD. After one day of incubation at 30 °C colony PCR and then sanger sequencing was performed to find frameshift mutation gene knockdowns.

### 4.4. sgRNA library design

Custom MATLAB scripts were used to design the optimized Cas9 library, and the key elements of the design are reported here. The optimized library had 3 guides designed for all 7919 mRNA coding genes in the *Y. lipolytica* CLIB89 genome (https://www.ncbi.nlm.nih.gov/assembly/GCA_001761485.1) (Magnan et al., 2016). Of these 3 guides, 2 were intended to be picked from the pool of best performing guides in the previous Cas9 screen (Schwartz et al., 2019a), while the third guide was designed by DeepGuide predictions (Baisya et al., 2022; Schwartz et al., 2019a). First, sgRNAs with CS>4.0 from the Schwartz et al. Cas9 screen were sorted from highest to lowest and the best two sgRNAs for each gene were identified. The third sgRNA for all genes, as well as guides for any genes that did not have two highly active guides (CS>4.0), were obtained from DeepGuide’s best predictions for that gene. All sgRNAs in the optimized library were verified to contain a unique seed sequence (11 nucleotides closest to the PAM). 360 nontargeting sgRNAs were also included in the library. These guides were confirmed not to target anywhere within the genome by ensuring that the first 12 nucleotides of the sgRNA did not map to any genomic loci.

### 4.5. sgRNA library cloning

The Cas9 library targeting the protein-coding genes in PO1f was ordered as an oligonucleotide pool from Agilent Technologies Inc. and cloned in-house using the Agilent SureVector CRISPR (Part Number G7556A). The library was subject to a NextSeq run to test for fold coverage of individual sgRNAs and skew. Cloned DNA was transformed into NEB 10-beta *E. coli* and plated. Sufficient electroporations were performed for each library to yield >100X library coverage. The plasmid library was isolated from the transformed cells after a short outgrowth. The optimized Cas9 library was cloned by making use of the Agilent SureVector CRISPR Library Cloning Kit (Part Number G7556A). Briefly, the backbone pCas9yl-GW was linearized and amplified by PCR using the primers InversePCRCas9Opt-F and InversePCRCas9Opt-R. To verify the completely linearized vector, we DpnI digested amplicon, purified the product with Beckman AMPure XP SPRI beads, and transformed it into *E. coli* TOP10 cells. A lack of colonies indicated a lack of contamination from the intact backbone. Library ssDNA oligos were then amplified by PCR using the primers OLS-F and OLS-R for 15 cycles as per vendor instructions using Q5 high fidelity polymerase. The amplicons were cleaned using the AMPure XP beads prior to use in the following step. sgRNA library cloning was conducted in four replicate tubes and subsequently, pooled and cleaned up as per manufacturer’s instructions.

One amplification bottle containing 1L of LB media and 3 g of high-grade low-gelling agarose was prepared, autoclaved, and cooled to 37 °C (Agilent, Catalog #5190-9527). Ten transformations of the cloned library were conducted using Agilent’s ElectroTen-Blue cells (Catalog #200159) via electroporation (0.2 cm cuvette, 2.5 kV, 1 pulse). Cells were recovered and with a 1 hr outgrowth in SOC media at 37 °C (2% tryptone, 0.5% yeast extract, 10 mM NaCl, 2.5 mM KCl, 10 mM MgCl2, 10 mM MgSO4, and 20 mM glucose.) The transformed *E. coli* cells were then inoculated into the amplification bottle and grown for two days until colonies were visible in the matrix. Colonies were recovered by centrifugation and subject to a second amplification step by inoculating two 250 mL LB cultures. After 4 hr, the cells were collected, and the pooled plasmid library was isolated using the ZymoPURE II Plasmid Gigaprep Kit (Catalog #D4202) yielding ∼1.8 mg of plasmid DNA encoding the optimized Cas9 sgRNA library. The library was subject to a NextSeq run to test for fold coverage of individual sgRNAs and skew.

### 4.6. *Y. lipolytica* library transformation

Transformation of the sgRNA plasmid libraries into PO1f and PO1f Cas9::A08 cells was accomplished with the method described in Chen et al, 1997, with modifications (Chen et al., 1997). For each strain of interest, seven 14 mL culture tubes, all containing 2 mL of YPD, were each inoculated with a single colony of the given strain, and grown in a 14 mL tube at 30 °C with shaking at 225 RPM for 22-24 hours (final OD ∼30). The cultures were then pooled, and 750 µL aliquots were distributed into 12 1.5 mL centrifuge tubes. Cells were pelleted (centrifuged at 4000 g), washed with ultrapure water, re-pelleted, and each resuspended in the Chen transformation buffer (53% PEG 4000 w/v, 0.1 M Lithium Acetate, 0.1 M DTT). Each tube was then sequentially dosed with 3 µL carrier DNA (ThermoFisher salmon sperm DNA, sheared, 10 mg/mL) and library plasmid stock (1 µg of plasmid per tube). Tubes were gently mixed and transformed by heat shock at 39°C for 1 hour. Each tube was recovered by adding 1 mL of fresh water, pelleting and aspirating supernatant. All 12 pellets were resuspended in 1 mL of fresh water, and then pooled back together. Dilutions of the transformation (0.1%, 0.01%, and 0.001%) were plated on solid SD-leu media to calculate transformation efficiency. The remaining volume of pooled transformants were then inoculated in 500 mL of SD-leu media (in a 2 L baffled flask), and grown for 48 hours at 225 RPM and 30 °C. Cells were centrifuged and resuspended in fresh SD-leu media to bring their OD to 20. Glycerol stock aliquots of the full volume of transformants were then prepared by mixing 800 µL of the resuspended transformation culture with 200 µL 100% glycerol, flash freezing in liquid nitrogen, and storing at −80 °C. Four biological replicates of each strain’s transformation were performed for pooling as necessary during screen experiments to ensure adequate diversity to maintain library representation and minimize the effect of plasmid instability (>100x coverage, >2.4 × 10^6^ total transformants per biological replicate).

### 4.7. *Y. lipolytica* acetate screen

CRISPR-Cas9 growth screens in acetate were conducted in synthetic defined minimal media from Turki et al., 2009 deficient in leucine (Turki et al., 2009). Three media conditions were prepared: Turki-leu 2% (w/v) glucose, Turki-leu 250 mM acetate, and Turki-leu 500 mM acetate. The pH of all media was adjusted to 6.3. Acetate was added as a sodium salt. 150 uL (approximately 1×10^7^ cells) of 2 day outgrowth glycerol stocks of PO1f Cas9 and PO1f strains transformed with the sgRNA library were used to inoculate 250 mL baffled flasks containing 25 mL of media. Two biological replicates were cultured for each different media and strain condition combinations. Outgrowth following inoculation was done at 30 °C at 220 RPM. The experiment was halted after 4 days of growth, where the OD_600_ of all flasks reached 8-12. On the last day, 1 mL of culture was removed, treated with DNase I, pelleted, and frozen for later library isolation and sequencing.

### 4.8. Acetate screen library isolation and sequencing

Frozen 1 mL DNase I treated and pelleted culture samples from each CRISPR screen flask was thawed and resuspended in 400 µL sterile Milli-Q water. Each cell suspension was split into two, 200 µL samples. Plasmids were isolated from each sample using a Zymo Yeast Plasmid Miniprep Kit (Zymo Research). Splitting into separate samples here was done to accommodate the capacity of the Yeast Miniprep Kit, specifically to ensure complete lysis of cells using Zymolyase and lysis buffer. This step is critical in ensuring sufficient plasmid recovery and library coverage for downstream sequencing as the gRNA plasmid is a low copy number plasmid. The split samples from a single pellet were pooled, and the plasmid number was quantified using quantitative PCR with qPCR-GW-F and qPCR-GW-R and SsoAdvanced Universal SYBR Green Supermix (Biorad). Each pooled sample was confirmed to contain at least 10^7^ plasmids so that sufficient coverage of the sgRNA library is ensured.

At least 0.2 ng of plasmids (approximately 3×10^7^ plasmid molecules) were used as template for PCR and amplified for 16 cycles and not allowed to proceed to completion to avoid amplification bias. The PCR product was purified using SPRI beads and tested on a bioanalyzer to ensure the correct length.

Samples from the screens were prepared as previously described in Schwartz et al., 2019. Briefly, isolated plasmids were amplified using forward (Cr1665-Cr1668) and reverse primers (Cr1669-Cr1673; Cr1709-1711) containing the necessary barcodes, pseudo-barcodes, and adapters (**Table S3**). Approximately 1×10^7^ plasmids were used as a template and amplified for 22 cycles, not allowing the reaction to proceed to completion. Amplicons at 250 bp were then gel extracted and tested on the bioanalyzer to ensure correct length. Samples were pooled in equimolar amounts and submitted for sequencing on a NextSeq 500 at the UCR IIGB core facility. All fitness scores (FS) for the acetate screens are provided in **Supp. File 1**.

### 4.9. Generating sgRNA read counts from raw reads

Next-generation sequencing raw FASTQ files were processed using the Galaxy platform (Afgan et al., 2018). Read quality was assessed using FastQC v0.11.8, demultiplexed using Cutadapt v1.16.6, and truncated to only contain the sgRNA using Trimmomatic v0.38. Custom MATLAB scripts were written to determine counts for each sgRNA in the library using Bowtie alignment (Bowtie2 v2..4.2; inexact matching) and naïve exact matching (NEM). The final count for each sgRNA was taken as the maximum of the two methods. Parameters used for each of the tools implemented on Galaxy are provided in **Table S5**, and the demultiplexing primers/barcodes are provided in **Supp. File 3**.

### 4.10. Identification of screening hits

Read counts from the glucose screen in SD-leu media in strains containing *KU70* gene knockout and Cas9 were used to calculate cutting score (CS) for each sgRNA by computing log2 ratio of the total normalized abundances in control and treatment samples, as described in (Ramesh et al., 2023). Similarly, read counts from glucose and acetate screens in Turki media were used to determine sgRNA fitness scores (FS) as log2 ratio of total normalized guide abundance in Cas9-containing treatment sample to that in the wildtype control. For the hydrocarbon screens, read counts from the Cas9-containing strain before inoculation onto solid media served as control to calculate sgRNA FS values. The sgRNA FS values from all datasets, along with the CS, were further used to identify significant hits from the respective screens via calculation of gene fitness scores using acCRISPR v1.0.0 (Ramesh et al., 2023). A CS-threshold of 1.5 was found to be optimum to remove low-activity sgRNAs from the optimized sgRNA library (library v2), and hence, the parameter *cutoff* was set to 1.5. Significance testing in acCRISPR is accomplished using a one-tailed or two-tailed z-test of significance. To identify essential genes for growth in glucose (Turki media), the parameter for significance testing (*significance*) was set to ‘one-tailed’ since only genes with statistically significantly lower fitness scores were to be called. To identify genes beneficial and detrimental to growth in the two acetate conditions as well as all hydrocarbon conditions, the *significance* parameter was set to ‘two-tailed’. In all cases, genes having FDR-corrected p-value < 0.05 were deemed as significant.

### 4.11. Hydrocarbon and Fatty Acid Solid Media Preparation

Solid medium hydrocarbon plates were prepared in batches of 20 (20 mL of media per 100 mm petri dish). First, 500 mL bottles of 2x concentration low melt agarose minimal media were prepared via autoclaving (for each bottle, 1 g MgSO_4_ * 7H_2_O, 4 g (NH_4_)_2_, 10 mg FeCl_3_ * 6H_2_O, 15 g LMP Agarose, 200 mg Uracil, 4 µg Myo-Inositol, 8 µg Biotin, 200 µg Thiamine HCl, 15 g KH_2_PO_4_, and 5.5 g K_2_HPO_4_), and kept molten in a 70°C water bath until ready for use. Meanwhile, for each plate type, bottles of 200 mL ultrapure water were autoclaved and kept heated to approximately 70°C on a heated stir plate. The amount of hydrocarbon or fatty acid used was calculated to bring the plates to the same molar amount of carbon provided by a 20 g/L glucose formulation, or 5.04 mL of Dodecane, 4.93 mL of 1-Dodecene 4.24 g of margaric (heptadecanoic) acid, and 4.67 mL of oleic acid. Hydrocarbon or fatty acid was directly added to the water, briefly allowed to mix (and melt if solid), and sonicated for 10 minutes (85% amplitude, 3 second on 3 second off cycle). Then, 200 mL of the 2x molten minimal media low melt agarose mix was added to the homogenized fatty acid/hydrocarbon water, briefly mixed on a stir plate, and immediately dispensed into petri dishes. Plates were allowed to harden in a laminar air flow hood for 2 hours and then refrigerated until needed, up to 3 days.

### 4.12. *Y. lipolytica* Hydrocarbon and Fatty Acid Solid Media Culturing

Optimized *Y. lipolytica* cell plating densities were calculated on solid medium hydrocarbon plates. Twelve hours prior to assay start, four glycerol stocks of prepared library cells (see methods 4.6), each from a separate transformation preparation, were pooled into 25 mL of SD-leu media in a 250 mL baffled flask, and grown for 12 hours at 225 RPM, 30°C. Cells were recovered as a pellet via centrifugation (at 4000 g for 5 minutes). To remove residual glucose, the cell pellet was washed with 15 mL of ultrapure water four times, with cells being recovered via centrifugation each time. Dilutions of the washed cells were then prepared to OD_600_ absorbances of 0.6, 0.4, 0.2, and 0.1. For each plate formulation to be tested, 100 µL of each cell dilution was plated on a single plate using sterile beads (4 plates per formulation). Plated cells were allowed to dry in a laminar air flow hood for 1 hour, then they were grown in a static 30°C incubator for up to 3 days. Starting at 24 hours, and every 24 hours thereafter, plates were checked for colony formation. Once discrete colonies had grown large enough to be easily visualized, the plates were removed and imaged in a Biorad imager. For each plate formulation, the day at which discrete colonies could be visualized was chosen as the endpoint for the selection screen assays. Finally, we selected the maximum OD for each plate formulation that still formed discrete (instead of fused) colonies to use as the selection screen plating density.

### 4.13. *Y. lipolytica* Hydrocarbon and Fatty Acid Plate Screen

Duplicate starter cultures of library cells were prepared 12 hours prior to the start of the assay as in section 4.12. Optimal optical density for plating on each hydrocarbon were calculated to be: OD600 0.1 for Oleic acid and OD600 0.2 for Dodecane, 1-Dodecene, and Margaric acid). For each plate formulation replicate, 100 µL of the chosen cell dilution was plated onto 8 pre-prepared plates (see section 4.11) using sterile beads (16 total plates per formulation). Plated cells were allowed to dry in a laminar air flow hood for 1 hour, then they were grown in a static 30°C incubator until the previously determined screen endpoint (see section 4.12; 2 days for Dodecane, 1-Dodecene, and oleic acid, and 3 days for Margaric acid). Upon removal from the incubator, plates were imaged in the BioRad imager.

The cells were then recovered in a two-part process prior to lysis. First, as much of the colony mass on the plate surface as possible was removed from each plate with 15 mL of dilute Tween 80 buffer (0.01% Tween 80, 0.11 M KH_2_PO_4_, 0.03 M K_2_HPO_4_) per plate, and the colonies scraped off with a plastic cell spreader. The samples from all eight plates were pooled for each replicate and centrifuged at 4000 g for 10 minutes to recover the cells. To recover the remaining cell mass which had burrowed into the solid media, we used a spatula to remove the solid media out of the plates, cut into small pieces, and placed in 250 mL centrifuge vessels. We added 35 g of urea to each vessel, and topped off to 100 mL with Tween 80 buffer. We then sealed and moved the vessels to a 70°C water bath for 10 minutes to melt the media and free the cell mass. The vessels were then centrifuged at 4000g for 5 minutes. The molten agar media was aspirated away from the cell pellets, which were rinsed in ultrapure water and re-centrifuged to remove any residual agar contamination, before being combined with the previously recovered surface colony cells. After pooling washed and embedded cells, we divided the total cell mass from each replicate into three 1.5 mL centrifuge tubes, and stored in a −80°C freezer until ready to harvest the library plasmids. The plasmid harvesting, quantification, barcode amplification, and sequencing steps were carried out in a similar manner as the acetate screen cells (see section 4.8 and 4.9), except no DNase I was used (as the harvesting process for the gel-embedded cell mass could have prematurely weakened the cell membranes and cultures were thoroughly washed). All fitness scores (FS) for the hydrocarbon screens are provided in **Supp. File 1**.

## Data availability

The sgRNA raw counts, cutting scores, and fitness scores generated in this study are provided as separate Supplementary Information and Source Data files. The sgRNA library will be made available at Addgene. The sgRNA sequencing data for the CRISPR-Cas9 screen generated for this study have been deposited in the NCBI SRA database under accession code PRJNA1125018.

## Author contributions

AR, VT, and IW conceived, planned, and analyzed data for the optimized gRNA library. NR, RJ, and MHD conceived and planned the acetate screen. BL and IW conceived and planned the hydrocarbon screens. NR conducted the acetate screen. BL conducted the hydrocarbon screens. NR, VT, AR, BL, and IW analyzed data. NR, YA, SL, AA, and CL conducted knockouts and growth experiments. VT and AN conducted acCRISPR analysis. NR, VT, BL, and IW wrote the manuscript. All authors edited the manuscript.

## Acknowledgments

This work was supported by DOE DE-SC0019093, NSF-2225878, NSF-1922642, and NSF-1803630.

