## Supplementary Material for "Optimized genome-wide CRISPR screening enables rapid engineering of growth-based phenotypes in *Yarrowia lipolytica*"

Table S1. Strains

Table S2. Plasmids

Table S3. Primers

Table S4. Guide RNAs

Table S5. Parameters for bioinformatics tools on Galaxy used in the analysis of NGS reads

Figure S1. Library v2 Characterization

Figure S2. Top Acetate Hit Characterization

Figure S3. Characterization of Top Validated Acetate Hits

**Table S1: Strains**

| **Strain** | **Description** | **Source** |
| --- | --- | --- |
| E. coli TOP 10 | F- mcrA Δ (mrr-hsdRMS-mcrBC) Φ80lacZΔM15 Δ lacX74 recA1 araD139 Δ (araleu)7697 galU galK rpsL (StrR) endA1 nupG | Thermo Fisher Scientific |
| Ys363 | Yarrowia lipolytica PO1f (URA3 KO, LEU2 KO) | ATCC |
| Ys1084 | Ys363 UAS1B8-TEF(136)-Cas9::A08 | Schwartz et al., 2019 |
| Ys1095 | Ys363 UAS1B8-TEF(136)-Cas9::A08, ku70 KO | Schwartz et al., 2019 |
| sNR004 | YS626 D05956g KO | This work |
| sNR005 | YS626 E22029g KO | This work |
| sNR006 | YS626 E13627g KO | This work |
| sNR007 | YS626 C02904g KO | This work |
| sNR008 | YS626 E01193g KO | This work |
| sNR009 | YS626 C33332g KO | This work |
| sNR010 | YS626 E24428g KO | This work |
| sNR011 | YS626 C30884g KO | This work |
| sNR012 | YS626 E37234g KO | This work |
| sNR013 | YS626 F23331g KO | This work |
| sNR014 | YS626 D01350g KO | This work |
| sNR015 | YS626 D21022g KO | This work |
| sNR016 | YS626 C23902g KO | This work |
| sNR017 | YS626 D12493g KO | This work |
| sNR018 | YS626 F11871g KO | This work |
| sNR019 | YS626 F19899g KO | This work |
| sNR022 | YS626 C02904g, E01193g KO | This work |
| sNR023 | YS626 C02904g, C33332g KO | This work |
| sNR024 | YS626 C02904g, E24428g KO | This work |
| sNR025 | YS626 C02904g, E37234g KO | This work |
| sNR026 | YS626 C02904g, D21022g KO | This work |
| sNR027 | YS626 E01193g, C33332g KO | This work |
| sNR028 | YS626 E01193g, E24428g KO | This work |
| sNR029 | YS626 E01193g, E37234g KO | This work |
| sNR030 | YS626 E01193g, D21022g KO | This work |
| sNR031 | YS626 C33332g, E24428g KO | This work |
| sNR032 | YS626 C33332g, E37234g KO | This work |
| sNR033 | YS626 E24428g, E37234g KO | This work |
| sNR034 | YS626 E24428g, D21022g KO | This work |
| sNR035 | YS626 E37234g, D21022g KO | This work |

**Table S2: Plasmids**

| **Plasmid** | **Description** | **Source** |
| --- | --- | --- |
| pIW715 | Homology donor plasmid for GFP integration at A08 (URA) | Addgene #84615 |
| pIW1009 | Homology donor plasmid for Cas9 integration at A08 (URA) | This work |
| pIW524 | Cas9 and sgRNA to cut at A08 (LEU) | This work |
| pIW386 | Easyclone Y. lipolytica Cas9 cutter without gRNA (LEU) | This work |
| pIW363 | pIW386 with gRNA for ku70 knockdown (LEU) | This work |
| pSC012 | pIW386 with URA instead of LEU marker | This work |
| pNR035 | D05956g KO gRNAin pSC012 | This work |
| pNR036 | E22029g KO gRNAin pSC012 | This work |
| pNR037 | E13627g KO gRNAin pSC012 | This work |
| pNR038 | C02904g KO gRNAin pSC012 | This work |
| pNR039 | E01193g KO gRNAin pSC012 | This work |
| pNR040 | C33332g KO gRNAin pSC012 | This work |
| pNR041 | E24428g KO gRNAin pSC012 | This work |
| pNR042 | C30884g KO gRNAin pSC012 | This work |
| pNR043 | E37234g KO gRNAin pSC012 | This work |
| pNR044 | F23331g KO gRNAin pSC012 | This work |
| pNR045 | D01350g KO gRNAin pSC012 | This work |
| pNR046 | D21022g KO gRNAin pSC012 | This work |
| pNR047 | C23902g KO gRNAin pSC012 | This work |
| pNR048 | D12493g KO gRNAin pSC012 | This work |
| pNR049 | F11871g KO gRNAin pSC012 | This work |
| pNR050 | F19899g KO gRNAin pSC012 | This work |

**Table S3: Primers**

| **Primer** | **Sequence** | **Description** |
| --- | --- | --- |
| A_FOR | CCACCGAGCCGTTCATG | D05956g forward primer |
| B_FOR | GTCCTGGAACGCCATCAG | E22029g forward primer |
| C_FOR | GCGTCTGAAGCTCGCTAATATCG | D16431g forward primer |
| D_FOR | CCGAGTGTAGGCCACTTG | E13627g forward primer |
| E_FOR | GTTAGACAGCACCAGGGTG | C02904g forward primer |
| F_FOR | GTTCTTGTCGTTGCAGACTCG | E01193g forward primer |
| G_FOR | CACTCACTTGCCACTGCAG | C33332g forward primer |
| H_FOR | CGTGATGGAGACTGGGGAG | E24428g forward primer |
| I_FOR | CCCGACTCTTCGTCTTCATCG | C30884g forward primer |
| J_FOR | GGTATGAATTCTGGCCCAAACTG | E37234g forward primer |
| K_FOR | GGCCTTCTCAGACAAGTCGG | F23331g forward primer |
| L_FOR | GTGATTGGGGTGTTAGGTCG | D01350g forward primer |
| M_FOR | GCATAAGTCTCAGAGCCAGC | D21022g forward primer |
| A_REV | CCAGACACACACACAGCC | D05956g reverse primer |
| B_REV | GCAACAACTTCCGAACTGCTG | E22029g reverse primer |
| C_REV | CCGTGTGGGACAATCTCTTTTAC | D16431g reverse primer |
| D_REV | GGAGAGAAAGACAGCGCTTTGC | E13627g reverse primer |
| E_REV | CTTTCCTCCAGACTTTTCTCCTTCC | C02904g reverse primer |
| F_REV | GGTTGGAGGGAATCGCG | E01193g reverse primer |
| G_REV | GTGGTAGTGGGCGAACTG | C33332g reverse primer |
| H_REV | CCTCCTACATTGCGCATGG | E24428g reverse primer |
| I_REV | CGAGTCGGCACTGAAGG | C30884g reverse primer |
| J_REV | GCATCTTGTGTCTGTAGAACCG | E37234g reverse primer |
| K_REV | GCTCACAGACACCTCTTGTG | F23331g reverse primer |
| L_REV | CGTTCGTCTGCACACACC | D01350g reverse primer |
| M_REV | CAACGACGTTGGGTGGC | D21022g reverse primer |
| neg1FOR | GATCAGAGTCAGGATGGGTGAC | C23902g forward primer |
| neg2FOR | GATAACGCCGTTCCACGC | D17181g forward primer |
| neg3FOR | CCTCGTTGCGATCCATTAACC | D12493g forward primer |
| neg4FOR | GTTGAGCTCATCCAGCTCATG | F21690g forward primer |
| neg5FOR | CTAATGCTGTCAAAACGGATAGCG | B06015g forward primer |
| neg6FOR | GCACCATGGTGGTAGGTG | B26350g forward primer |
| neg7FOR | GCTCAATCCACCACAAGATCAAATC | F11871g forward primer |
| neg8FOR | CGAGGCATTACCTTTGGAGG | D11769g forward primer |
| neg9FOR | CAGCAGTAGCCCCAACAC | A22093g forward primer |
| neg10FOR | CTCCACACTTGGCAGTGG | A21156g forward primer |
| neg11FOR | gcACTCGTCTTTGAGTCTCAC | F07819g forward primer |
| neg12FOR | CCAGAACCGTTATTGGTGCC | B05364g forward primer |
| neg13FOR | CGTCACAGGTATCGGCAAC | D14893g forward primer |
| neg14FOR | gatcttcttgccGTCGGC | D15424g forward primer |
| neg15FOR | gtggaagaggttgTGGCC | A21797g forward primer |
| neg16FOR | GTCTTCCTTGTCCACCCG | F34749g forward primer |
| neg17FOR | CCCAACCTGTATTTCGGTGTC | F19899g forward primer |
| neg18FOR | GGGTATGAGAGGAATTTCGacg | A03183g forward primer |
| neg19FOR | GTACTGGCAGAAGAGCGC | D25330g forward primer |
| neg20FOR | CATCTTACCACCCTGAGTAAGTCC | C21487g forward primer |
| neg21FOR | GAGGCATCCGGTTCGTTC | F26726g forward primer |
| neg1REV | GCAAATCAATTCTTCGTGCACG | C23902g reverse primer |
| neg2REV | GTTGTAGATTGTCAGGACCATAATGG | D17181g reverse primer |
| neg3REV | CACCTGTTCCATGAGCGC | D12493g reverse primer |
| neg4REV | GAAGGATGTCTAGATGAACCACG | F21690g reverse primer |
| neg5REV | GATATCTCCGAGACGAGTTTTCCTC | B06015g reverse primer |
| neg6REV | CAACACTGACGTGACGTCC | B26350g reverse primer |
| neg7REV | CCACCAAGCTGCTTGGATAAG | F11871g reverse primer |
| neg8REV | GCTGAGTGTTAGAACAACTCGC | D11769g reverse primer |
| neg9REV | CACCAAAGAACTGTTTGGAGGG | A22093g reverse primer |
| neg10REV | CCGAGACCGAATCCTCGag | A21156g reverse primer |
| neg11REV | GCGAAGAAACCAGTGCCTG | F07819g reverse primer |
| neg12REV | GGAACTTAGTAGCCTCGATAGGTC | B05364g reverse primer |
| neg13REV | CGTTGTCGACGAGGAATcc | D14893g reverse primer |
| neg14REV | GTCTGGTGATGTTGTCTTGGTG | D15424g reverse primer |
| neg15REV | GTTGGTCTTAGTGCTGACGC | A21797g reverse primer |
| neg16REV | CAGTCATTCGCTTTCGAGATGAAG | F34749g reverse primer |
| neg17REV | CAATTCGCTCCTGCAGAAGG | F19899g reverse primer |
| neg18REV | GACGTTGAACACGGAAATGCC | A03183g reverse primer |
| neg19REV | CAGCTCCAACGGGATATTGC | D25330g reverse primer |
| neg20REV | CGTAACCTCCAGGAGGTTCTC | C21487g reverse primer |
| neg21REV | GGTTGAACAGCGCGTCC | F26726g reverse primer |
| qPCR-GW-F | TTATGAACTGAAAGTTGATGGC | qPCR forward primer |
| qPCR-GW-R | TCACACAGGAAACAGCTATG | qPCR reverse primer |
| Cr_1665 | AATGATACGGCGACCACCGAGATCTACACTCTTTCCCTACACGACGCTCTTCCGATCTAGTCCGGTTCGATTCCGGGTC | Forward primer Illumina |
| Cr_1666 | AATGATACGGCGACCACCGAGATCTACACTCTTTCCCTACACGACGCTCTTCCGATCTGTAGTCCGGTTCGATTCCGGGTC | Forward primer Illumina |
| Cr_1667 | AATGATACGGCGACCACCGAGATCTACACTCTTTCCCTACACGACGCTCTTCCGATCTCAGTAGTCCGGTTCGATTCCGGGTC | Forward primer Illumina |
| Cr_1668 | AATGATACGGCGACCACCGAGATCTACACTCTTTCCCTACACGACGCTCTTCCGATCTTCCAGTAGTCCGGTTCGATTCCGGGTC | Forward primer Illumina |
| Cr_1669 | CAAGCAGAAGACGGCATACGAGATTCGCCTTGGTGACTGGAGTTCAGACGTGTGCTCTTCCGATCTCGACTCGGTGCCACTTTTTCAAG | Reverse primer Illumina |
| Cr_1670 | CAAGCAGAAGACGGCATACGAGATATAGCGTCGTGACTGGAGTTCAGACGTGTGCTCTTCCGATCTCGACTCGGTGCCACTTTTTCAAG | Reverse primer Illumina |
| Cr_1671 | CAAGCAGAAGACGGCATACGAGATGAAGAAGTGTGACTGGAGTTCAGACGTGTGCTCTTCCGATCTCGACTCGGTGCCACTTTTTCAAG | Reverse primer Illumina |
| Cr_1672 | CAAGCAGAAGACGGCATACGAGATATTCTAGGGTGACTGGAGTTCAGACGTGTGCTCTTCCGATCTCGACTCGGTGCCACTTTTTCAAG | Reverse primer Illumina |
| Cr_1673 | CAAGCAGAAGACGGCATACGAGATCGTTACCAGTGACTGGAGTTCAGACGTGTGCTCTTCCGATCTCGACTCGGTGCCACTTTTTCAAG | Reverse primer Illumina |
| Cr_1709 | CAAGCAGAAGACGGCATACGAGATGTCTGATGGTGACTGGAGTTCAGACGTGTGCTCTTCCGATCTCGACTCGGTGCCACTTTTTCAAG | Reverse primer Illumina |
| Cr_1710 | CAAGCAGAAGACGGCATACGAGATTTACGCACGTGACTGGAGTTCAGACGTGTGCTCTTCCGATCTCGACTCGGTGCCACTTTTTCAAG | Reverse primer Illumina |
| Cr_1711 | CAAGCAGAAGACGGCATACGAGATTTGAATAGGTGACTGGAGTTCAGACGTGTGCTCTTCCGATCTCGACTCGGTGCCACTTTTTCAAG | Reverse primer Illumina |

**Table S4: Guide RNAs**

| **Guide** | **Sequence** | **Description** |
| --- | --- | --- |
| YALI1_D05956g_2 | CAACATGTACGGCTTCCATA | D05956g guide |
| YALI1_E22029g_3 | CACTCCTCTGTTGAGTGCGG | E22029g guide |
| YALI1_D16431g_3 | AACCATGACAAATCTGCTCA | D16431g guide |
| YALI1_E13627g_3 | TGGATGTGTGTGACGGAAAG | E13627g guide |
| YALI1_C02904g_3 | AAAGCCGAAGATGGGGCCCG | C02904g guide |
| YALI1_E01193g_3 | CTCTCGGTTTCATTCCCGAG | E01193g guide |
| YALI1_C33332g_2 | CGGAAGATATGACGATAAAG | C33332g guide |
| YALI1_E24428g_3 | GTGTGTCTTGAGGTCCCGCT | E24428g guide |
| YALI1_C30884g_2 | GCAAGAGCCGAGTCAGCGCA | C30884g guide |
| YALI1_E37234g_1 | ATTGCAGGTTAGAAATGGGG | E37234g guide |
| YALI1_F23331g_2 | CCTTGAGCTTGAGACCCTGC | F23331g guide |
| YALI1_D01350g_1 | AGTTGTCAGTGGCAAGGTAG | D01350g guide |
| YALI1_D21022g_1 | GATGCTCCGACGAGGCCTGC | D21022g guide |
| YALI1_C23902g_1 | CGATGACTCTGGGCACGAGG | C23902g guide |
| YALI1_D17181g_1 | CTACTCGGTCTACAGACGAG | D17181g guide |
| YALI1_D12493g_3 | GGAGGGCGAAGAGAGGGTCG | D12493g guide |
| YALI1_F21690g_2 | GTACTCGGTGGACGACAGTT | F21690g guide |
| YALI1_B06015g_1 | TCAGCTAAACCCATGTCAAA | B06015g guide |
| YALI1_B26350g_3 | ATGATGGCAATGATCACGGG | B26350g guide |
| YALI1_F11871g_2 | CCACCGACTTGAGAATGCCT | F11871g guide |
| YALI1_D11769g_2 | GGTGTCATCAACCGACACTC | D11769g guide |
| YALI1_A22093g_3 | TCTCCCAGACAGGTCCAGAT | A22093g guide |
| YALI1_A21156g_1 | GTATCGTGCCTTAGGCCAGG | A21156g guide |
| YALI1_F07819g_3 | GAAACCCGCAAAAACCTCCA | F07819g guide |
| YALI1_B05364g_1 | TCAGAGCGTTAAGAATAGCG | B05364g guide |
| YALI1_D14893g_3 | CGCCTCCGTGTGCTACATCT | D14893g guide |
| YALI1_D15424g_3 | ATCCTTGATGGAGAGCTTGT | D15424g guide |
| YALI1_A21797g_1 | GCAGAGCCTGGAAGAACCCA | A21797g guide |
| YALI1_F34749g_3 | CCTGGGCGAGGAGAGCGATG | F34749g guide |
| YALI1_F19899g_1 | AAGACCAGAGTAGAACACCA | F19899g guide |
| YALI1_A03183g_1 | GAGCGACTAGGCACTCTCGA | A03183g guide |
| YALI1_D25330g_1 | GCAGATACGACAGCTCTGAG | D25330g guide |
| YALI1_C21487g_3 | TGTACTCGTAGTACTGCACT | C21487g guide |
| YALI1_F26726g_2 | ATATTGCACAAAGTGGACCA | F26726g guide |

**Table S5.** Parameters for bioinformatics tools on Galaxy used in the analysis of NGS reads

| Tool | Version | Parameters* |
| --- | --- | --- |
| FastQC | v0.11.8 | Default settings |
| Cutadapt | Galaxy Version 1.16.6 ^13^ | Cutadapt was used to demultiplex samples containing the same Illumina barcode, but different pseudobarcodes at the 5’ end of the read. Samples were amplified with reverse primers Cr1669-1673;Cr1709-1711 and forward primers Cr1665-1668 each containing a different pseudo barcode as mentioned in Table   - 5’ (Front) anchored 6 bp pseudo-barcodes to be demultiplexed (-g): ^NNNNNN (refer to previous table for pseudo-barcode-forward primer association). - Maximum error rate (--error-rate): 0.2 - Match times (--times): 1 - Minimum overlap length (--overlap): 4 - Multiple output: Yes (Each demultiplexed readset is written to a separate file) |
| Trimmomatic | v0.38 | - HEADCROP: 30 (if amplified by Cr1665); or 32 (if amplified by Cr1666); or 34 (if amplified by Cr1667); or 36 (if amplified by Cr1668) - CROP: 20 |
| Bowtie2** | v2.4.2 | - Number of allowed mismatches in seed alignment (-N): 1 - Length of the seed substring (-L): 20 - Function governing interval between seed substrings in multiseed alignment (-i): S,1,0.50 - Function governing maximum number of ambiguous characters (--n-ceil): L,0,0.15 - Alignment mode: end-to-end - Number of attempts of consecutive seed extension events (-D): 20 - Number of times re-seeding occurs for repetitive reads: 3 - Save mapping statistics: Yes |

* All parameters other than those mentioned here are kept at default values.

** Bowtie2 usage needs a genome fasta file for alignment. Nontargeting sgRNA and any other sgRNA that Bowtie2 could not find within the original CLIB89 genome file were appended as an extra chromosome so that Bowtie could align all sgRNA for the purposes of generating counts.

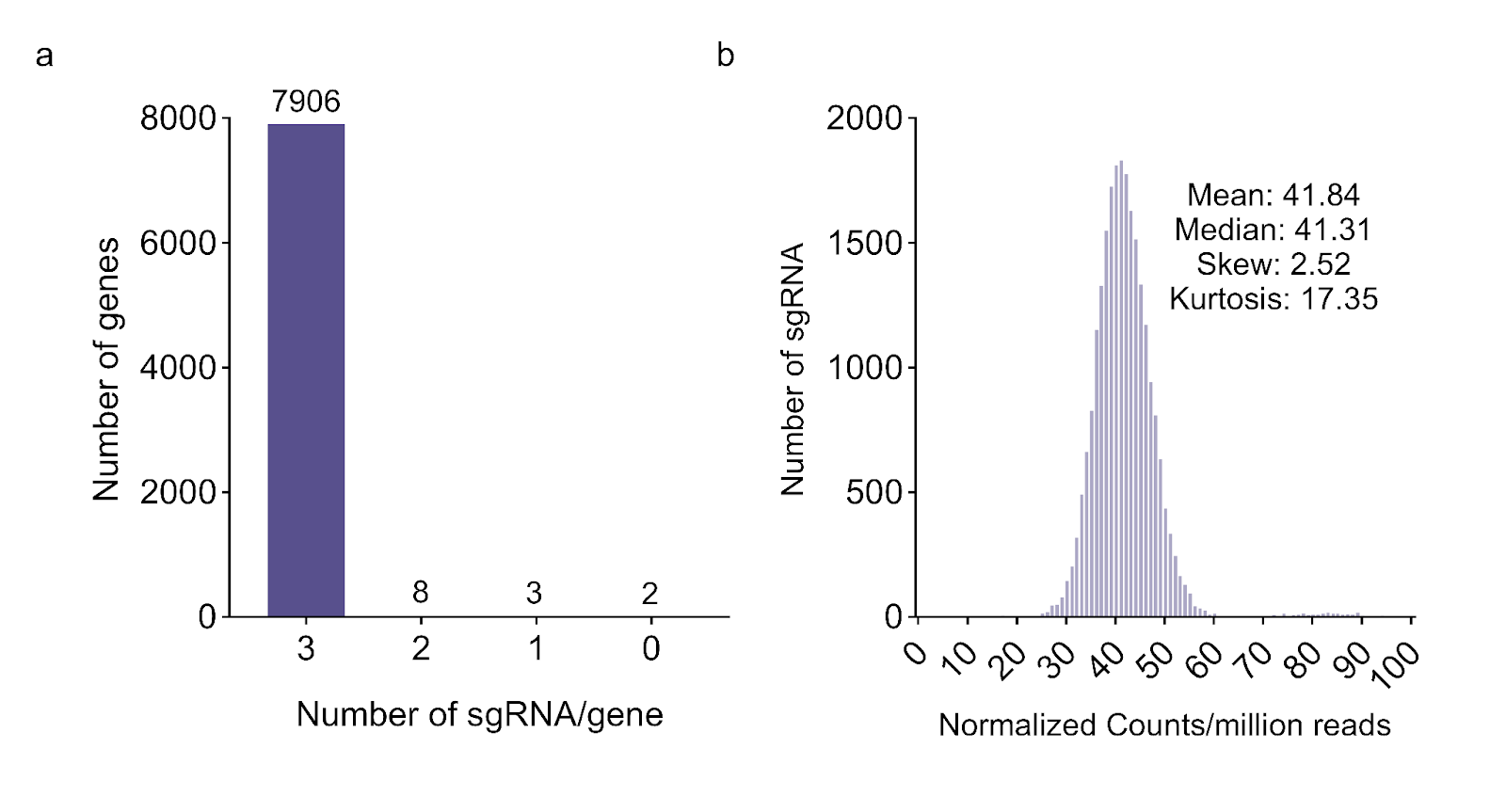

**Figure S1: Library v2 Characterization.** Characteristics of lib. v2. (a) 99.8% of genes have three targeting guides. (b) Untransformed lib. v2 contains a tight distribution of guides.

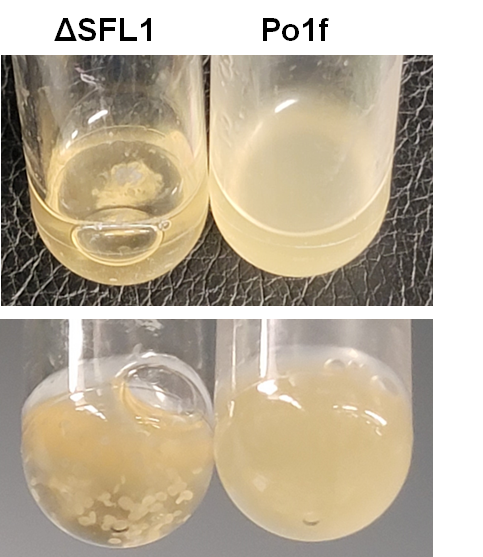

**Figure S2: Top Acetate Hit Characterization**. Top acetate positive FS hit encodes flocculation suppressing gene SFL1. The ΔSFL1 knockout causes *Y. lipolytica* to flocculate.

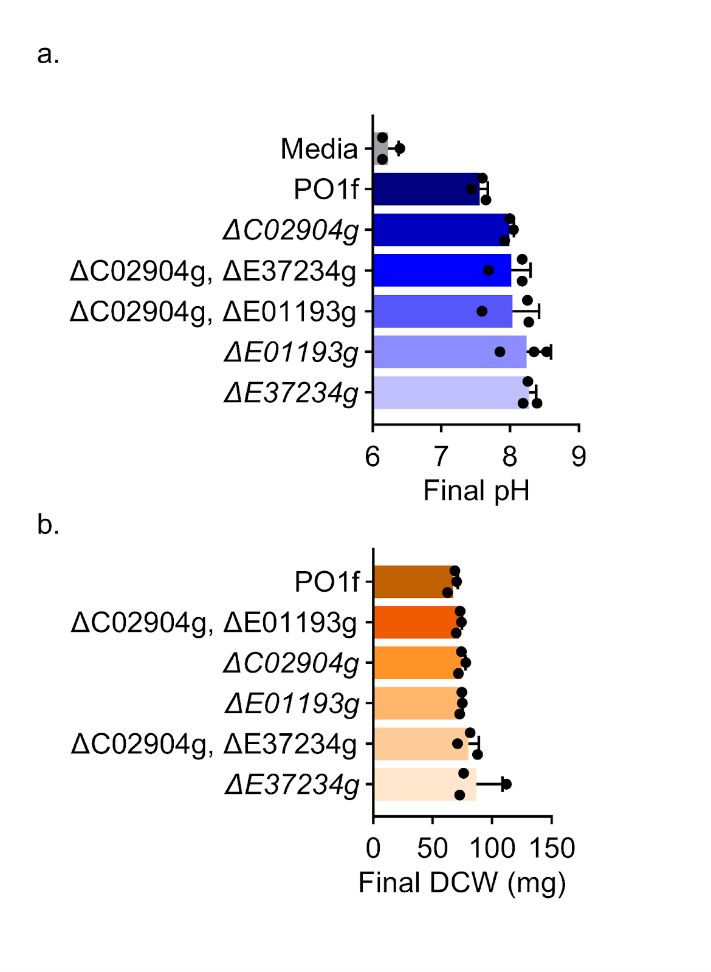

**Figure S3: Characterization of Top Validated Acetate Hits.** Top acetate positive FS hits were characterized at the end of a growth curve performed in 500 mM acetate minimal media in 250 mL shake flasks 30 ℃, 220 RPM shaking, 5 days. (a) Higher levels of growth cause the pH of the media to drop more as acetate is consumed. (b) 10 mL of cell culture was collected at the end of 5 days, pelleted, lyophilized, and weighed. Top positive FS hit knockouts grow more quickly and produce more biomass than the wild type.
